# Longitudinal Surveys Show that Urethral Catheters Harbor Recurrent Polymicrobial Biofilms with Cooperative and Competitive Behaviors Among Cohabitating Pathogens

**DOI:** 10.1101/388389

**Authors:** Yanbao Yu, Harinder Singh, Tamara Tsitrin, Shiferaw Bekele, Kelvin J. Moncera, Patricia Sikorski, Manolito G. Torralba, Lisa Morrow, Randall Wolcott, Karen E. Nelson, Rembert Pieper

**Affiliations:** J. Craig Venter Institute, 9605 Medical Center Drive, Rockville, Maryland 20850; J. Craig Venter Institute, 4120 Capricorn Lane, La Jolla, California 92037; Southwest Regional Wound Care Center, 2002 Oxford Avenue, Lubbock, Texas 79410

## Abstract

The analysis of catheter biofilms (CBs) from patients with neurogenic bladder disorders revealed persistent colonization by polymicrobial communities. The recurrence of bacterial species in the CBs of sequentially replaced catheters suggests urothelial reservoirs responsible for recolonization of the catheter surface. Proteomic data for CB samples were indicative of chronic inflammation in the patients’ urinary tracts via neutrophil and eosinophil infiltration and epithelial cell exfoliation. These host defense pathways, effective in killing pathogens during uncomplicated urinary tract infection, failed to eliminate CBs. Intermittent antibiotic drug treatment had different outcomes: either replacement of drug-susceptible by drug-tolerant bacteria or transient microbial biomass reduction followed by resurgence of the previously colonizing bacteria. Proteins that sequester iron and zinc such as lactotransferrin, lipocalin-2 and calprotectin were abundant in the patient’s CBs and urine sediments. Indicative of a host-pathogen battle for bivalent metal ions, acquisition and transport systems for such ions were highly expressed by many organisms residing in CBs. Species part of the Enterococcaceae and Aerococcaceae families, generally not well-characterized in their ability to synthesize siderophores, frequently cohabitated biofilms dominated by siderophore-producing *Enterobacteriaceae.* In support of metal acquisition cooperativity, we noticed positive abundance correlations for a *Proteus mirabilis* yersiniabactin-type siderophore system and two *Enterococcus faecalis* ABC transporters. Distinct bacterial strains highly expressed known or putative cytotoxins that appeared to compromise the survival of co-resident bacteria, e.g. a *P. mirabilis* hemolysin and *Pseudomonas aeruginosa* type 6 secretion and pyoverdin biosynthesis systems. In conclusion, there is support for cooperative and competitive behaviors among bacteria cohabitating CBs.

## Introduction

Urethral catheter-associated urinary tract infections (CAUTIs) are a common type of complicated UTI. CAUTIs have a higher risk of recurrence, pyelonephritis and bacteremia than uncomplicated UTIs in the nosocomial environment (1–3). Asymptomatic cases are usually diagnosed as catheter-associated asymptomatic bacteriuria (CAASB). The use of nearly 100 million urethral catheters per year worldwide, the 3% to 10% incidence of bacteriuria over 24 hours following patient catheterization and an average bladder catheter insertion time of 72 h (2) suggest an estimated 9 to 27 million CAUTIs per year globally. The most common microbial causes are *Escherichia coli, Klebsiella pneumoniae, Pseudomonas aeruginosa, Proteus mirabilis, Enterococcus* and *Candida spp.* (1, 3, 4). Indwelling Foley catheters are often used by patients with anatomical urinary tract abnormalities and neurogenic bladders and retained in the urinary tract for two or more weeks. Even if catheters are replaced, microbial colonization is known to be difficult to avoid without antibiotic treatments. Bacteria able to form mucoid biofilms *(e.g., E. faecalis, P. aeruginosa, K. pneumoniae,* and uropathogenic *E. coli)* and those degrading urea as a carbon source *(e.g., Proteus* and *Providencia spp.)* are dominant in polymicrobial communities forming on catheter surfaces (3, 5, 6). A degradation product of urea, ammonium, alkalinizes the pH of urine and triggers precipitation of phosphate salt crystals on the catheter surface, thus increasing the risk of luminal occlusion and complications such as urinary stones and kidney infections (3). Unless specific risk factors exist *(e.g.,* a compromised immune system or pregnancy), clinical guidelines do not recommend the use of antibiotics for CAASB (7). Many uropathogens have genetically acquired and innate resistances against several classes of antibiotic drugs. This includes the ESKAPE group of pathogens (8). Understanding the mechanisms involved in microbial cohabitation and competition in catheter surface biofilms may lead to new approaches to prevent or perturb their formation and renewal following antibiotic treatments.

The pathogenesis of UTI and CAUTI has been studied extensively in the context of the pathogens *E. coli* and *P. mirabilis* (4, 9, 10). Bacterial molecular pattern recognition by urothelial cell effectors and receptors triggers Toll-like receptor (TLR) signaling events which then result in leukocyte infiltration. Neutrophils are the immune cells primarily responsible for rapid phagocytosis of the invading pathogens (11). Bacterial clearance via neutrophil effector functions include the release of granular proteins and reactive oxygen species and the formation of extracellular traps (12, 13). Studies of *E. coli*-associated recurrent UTI have implicated the neutrophil cyclooxygenase-2 in the pathogenesis of recurrence (14). The latter appears to be influenced by host susceptibility and the urovirulence of strains colonizing the human intestinal tract (15). Type I fimbriae, expressed by several Gram-negative uropathogens, are thought to initiate an infection by binding to mannoslyated uroplakins, glycoproteins that coat the surface of urothelial umbrella cells (16). *Proteus mirabilis* is the major cause of encrusted catheter biofilms with salt deposition (3, 9). Viable bacterial cells disperse and recolonize the catheter. Their ascendance to the kidneys can enhance the disease severity (3). The signals that control microbial biofilm dynamics over time are complex and involve the availability of nutrients including various carbon sources and bivalent metal ions. Iron and zinc are sequestered by the immune system during infections (9, 17, 18). The urinary pH and exposure to antimicrobial effectors and antibiotic drugs affect disease outcome (2, 4). Mucoid biofilms are known to involve the deposition of host proteins on catheter surfaces to which bacteria can adhere (3, 4). Part of the mucus are polysaccharides that encapsulate the bacterial cells and lessen the effectiveness of innate immune responses. Like encrusted biofilms, mucoid biofilms can cause renal complications by blocking urine flow (3).

Microbial culture-based approaches have shown that urethral catheter biofilms are biologically complex (5, 6, 19). Culture-independent metagenomic surveys have identified fastidious and rarely pathogenic microbes, *e.g. Actinomyces, Stenotrophomonas, Corynebacterium, Anaerococcus, Achromobacter* and *Finegoldia spp.* (20, 21), suggesting more diversity in catheter biofilms than previously known. Little is known about the community dynamics and their fate upon sequential catheter exchanges in a patient. Additional unsolved questions pertain to the chronicity of host immune responses and the patterns in which biofilms challenged by antibiotic treatment survive. Bladder catheterization was found to cause persistent sterile inflammation with CD^45^+ neutrophils as the main infiltrating immune cells in a study of a murine CAUTI model. Increased susceptibility to *E. coli* and *E. faecalis* infections was accompanied by urothelial barrier disruption and further immune cell infiltration (22). Co-infection with *E. coli* and *P. mirabilis* in a murine UTI model suggested that these species utilize central carbon metabolites in a complementary fashion (23). Among our objectives was to use metaproteomic methods applied to patient specimens to understand colonization patterns over a series of urethral catheter biofilms from a distinct patient, looking at mutualistic as well as competitive behaviors.

## Results

### SCI patient cohort, clinical and phenotypic observations

The patients (2 females and 7 males; 4 hispanics and 5 caucasians) were adults with spinal cord injuries who suffered from neurogenic bladder syndrome. Chronically infected wounds were a comorbidity for all patients. During clinical visits when infected wounds were attended to, urethral catheters were typically replaced. Topical wound treatments were the norm, and only three patients received systemic antibiotic treatments. Dataset S1 (Suppl. Data) contains antibiotic treatment information along with other patient metadata. No patient reported symptoms indicative of UTI, consistent with the diagnosis of CAASB, at any time point of the survey. The catheters were replaced every 1 to 3 weeks to reduce the extent of bacteriuria and risk of renal complications. We analyzed UP and CB samples from serially collected catheter and urine specimens, longitudinally sampling over a time of two to six months with up to 15 timepoints. Visual inspection revealed evidence of salt-encrusted catheter biofilms for specimens from P4, P5, P6 and P7 (Table 1). Average biomasses of encrusted catheters were moderately larger those of mucoid biofilms. In some cases, the biofilm biomass was mostly dispersed into urine (UP samples). In other cases, the biomass was mostly found in CB samples indicative of stronger adherence.

**Table 1.** Biomass and visual appearance of catheter biofilms

|  | P1 (M) | P2 (F) | P3 (M) | P4 (M) | P5 (M) | P6 (M) | P7 (F) | P8 (M) | P9 (F) |
| --- | --- | --- | --- | --- | --- | --- | --- | --- | --- |
| wUP (g) | 0.12 | 0.07 | 0.03 | 0.27 | 0.18 | 0.23 | 0.40 | 0.05 | 0.25 |
| wCB (g) | 0.20 | 0.14 | 0.11 | 0.10 | 0.29 | 0.75 | 0.10 | 0.06 | 0.15 |
| Biofilm | mucoid | Mucoid | mucoid | salt-encr | salt-encr* | salt-encr | salt-encr | mucoid | mucoid |

P1 to P9: patient numbering; (M) male, (F) female; biomasses are provided in wet cell weight for ~ 30 ml urine (wUP) and estimated for the entire length of a catheter (wCB). Values represent averaged masses from analyses of 2 to 8 specimens per patient. \*In this case, salt encrustation (salt encr) was noted for 3 early timepoints (specimens) but not for 5 specimens collected later.

### Omics analysis overview

UP and CB_pel_ samples from eight of nine patients were subjected to 16S rDNA metagenomic analysis, yielding 112 datasets for bacterial genera. UP and CB samples from eight of nine patients were used for metaproteomic analysis to determine species or genus abundances and examine their proteomes. Metaproteomic data were also suitable to survey the immune responses against pathogens and urothelial tissue injury and repair events in patients. The 16S rDNA data also served to customize proteomic database contents in a patient-specific manner (see Methods). The comparison of search results using different species from the same genus allowed us to select the correct bacterial species for a series of samples from a patient under study. In the selection process, existing knowledge on microbial species that cause UTIs was taken into consideration. Using this trial-and-error and time-consuming approach, we were confident that all major colonizing microbial species were represented in the datasets. As the isolation of microbial colonies from *in vitro* cultures in some cases showed, species near or below the detection limit of proteomic analysis in clinical samples may have been missed. Given that we were interested in the overall dynamics of microbial colonization and the resilience of distinct strains or species on catheters over time, as well as the immune responses that were likely driven by the microbes more abundant in CBs, our approach did not diminish quality of results and conclusions. We generated 121 metaproteomic datasets that allowed us to investigate population complexity and dynamics in the catheter biofilm milieu in eight patients over several months and simultaneously yield insights into the chronicity of host immune responses. Dataset S2 (Suppl. Data) has all proteins, descriptions and normalized protein quantities.

### 16S rDNA phylogenetic profiles indicate persistent colonization of catheters by uropathogens and other bacteria adapted to life in the urogenital tract

To our knowledge, we show for the first time that combinations of bacterial genera colonize catheters and persist as biofilms despite frequent replacements of this medical device in the patients’ urinary tracts over up to six months. The compositional variation of such communities is low comparing UP and CB samples from the same timepoint, but often higher when comparing samples derived from different timepoints. Different patients have entirely different CB communities. 16S rDNA phylogenetic analyses revealed a total of 40 genera with OTU abundances greater than 0.07%, averaged from all 112 analyzed samples (Dataset S3, Suppl. Data). For distinct UP/CB sample timepoints, the number of identified genera ranged from 2 to 15. While Firmicutes and Proteobacteria resided in catheters from all patients, Actinobacteria, Bacteroides and Fusobacteria were present in fewer patients. Among the most prevalent genera were those that are known causes of CAUTI and UTI: *Proteus, Escherichia, Klebsiella, Citrobacter, Enterococcus* and *Staphylococcus*. Genera representative of species reported to rarely cause UTI but found in catheter biofilms were limited to one or two patients, such as *Bordetella, Aerococcus* and *Actinobaculum.* 16S rDNA data are quantitatively not accurate due to genus-specific differences in the PCR efficiency of the targeted V1-V3 region. We resorted to proteomics for quantitative data interpretations of microbes identified in CBs. As displayed in Fig. 1, the diversity of bacterial genera was lower in salt-encrusted biofilms (P4, P6, P7) than in mucoid biofilms (P1, P2, P8). Early P5 samples were placed in the former category, P5 samples collected later were in the latter group. The Shannon index accounts for both abundance and evenness of genera/OTUs. Our findings support the notion that the alkaline pH milieu in urine and salt crystallization on catheter surfaces favor growth of the Proteus/Providencia group of bacteria which produce ureases and are well-adapted to this milieu (24).

**Fig. 1.**
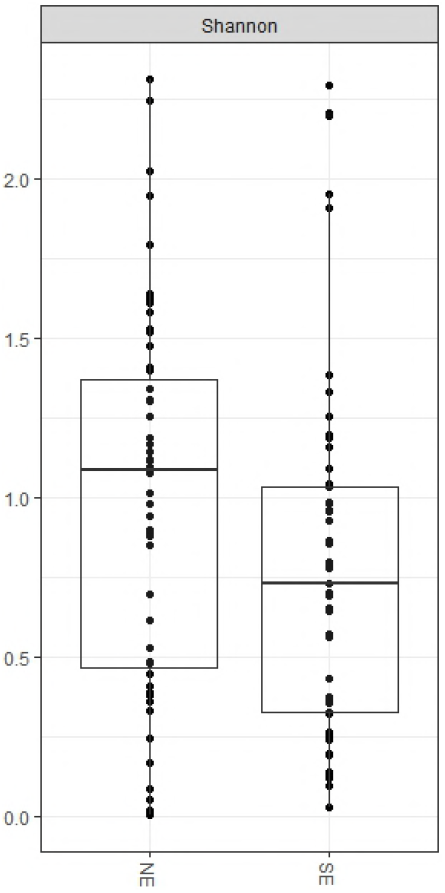
Diversity of bacterial genera using calculations based on the Shannon index of diversity. NE and SE: non-encrusted and salt-encrusted biofilms, respectively. The 16S rDNA data from UP and CB samples for a given time point were considered separately in this analysis.

### Proteomic data reveal dominance of *P. mirabilis* in salt-encrusted CBs joined by microbial species that vary more in the comparison of different patients than for specimens from a distinct patient

UP and CB samples were collected from P4, P5, P6 and P7 over 3.5, 6, 4 and 5.5 months, respectively. The proteomic data-based graphics in Fig. 2 show that *P. mirabilis* was dominant in nearly all samples derived from P4 and P6 (and P7, as presented in Fig. S1, Suppl. Data). In the samples of P5, *P. mirabilis* was displaced over time by its phylogenetic relatives *Morganella morganii* and *P. stuartii* and by *E. coli*. The UP/CB sample series for P4, P5 and P6 show that microbial species of the order Lactobacillales *(E. faecalis, Aerococcus urinae* and *Globicatella)* persisted over several catheter replacements but rarely out-competed *P. mirabilis* or another member of the Enterobacteriaceae. Clearly, these small cocci have adapted to life on catheter surfaces and thrive in community biofilms joined by various Gram-negative pathogens. In the CBs of P5 and P6, *P. stuartii* was moderately abundant, thus also suggesting benefits of cohabitation with other Gram-negative pathogens in a polybacterial biofilm. Noteworthy is the persistent presence of *Actinobaculum massiliense,* a rarely described actinobacterial species that has only once before been linked to chronic UTI (25), to the P5 datasets. We observed that microbial contributions to CB proteomes were typically higher than those to UP proteomes. Human protein content (with a subset shown for the four categories NEI, CAP, ERY and KRT) in UP proteomes was more than 75% of the total protein in most instances (Fig. 2), due to release of immune cells, gel-forming uromodulin, and urothelial and squamous epithelial cells shed during the inflammatory process. In summary, quantitative proteomic changes in sequentially collected samples reveal a dynamic interplay of bacterial species with evidence of adherence and recolonization of newly placed urethral catheters and biofilm dispersal. Host cell functional data displayed in Fig. 2 are characterized later in the Results section.

**Fig. 2.**
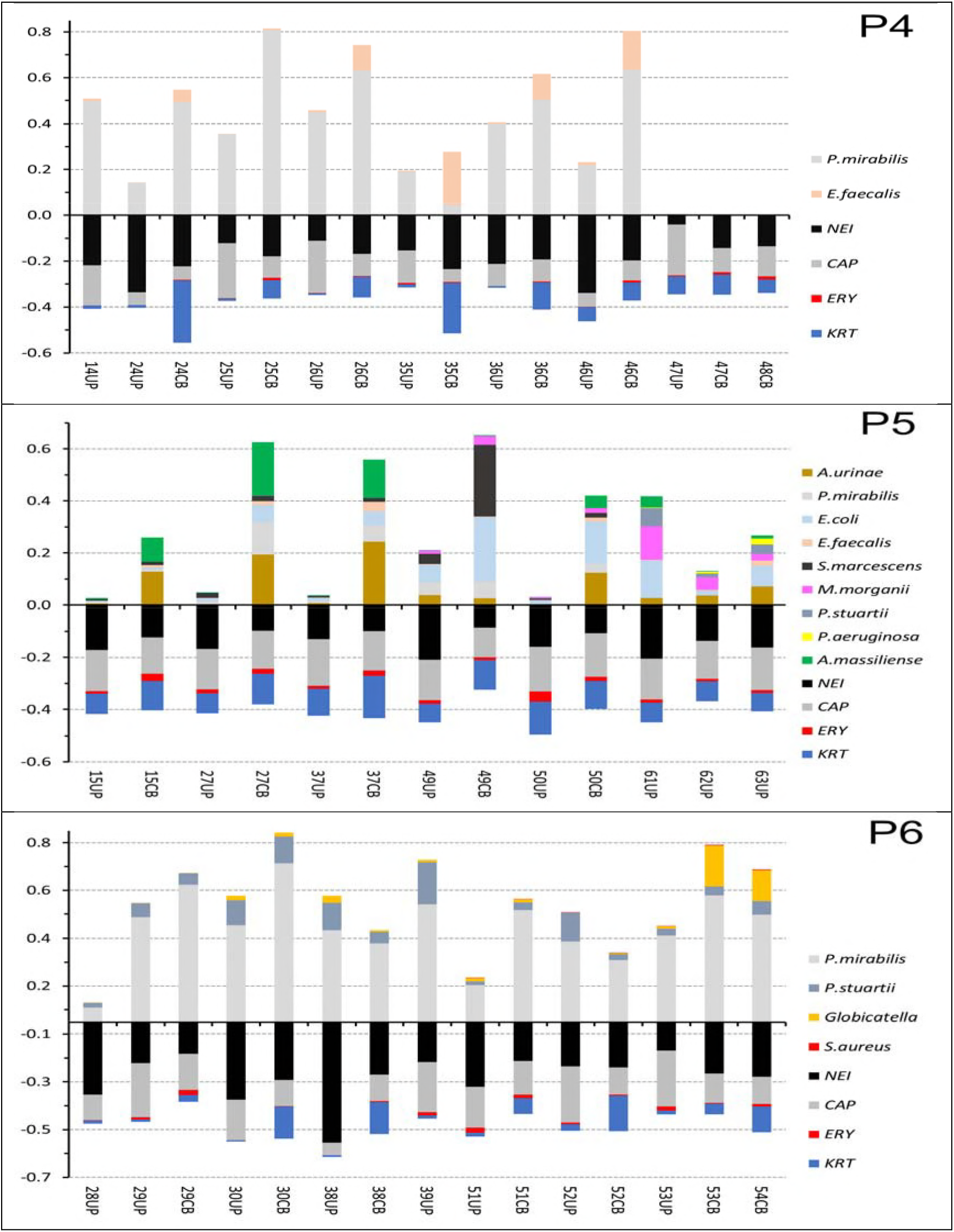
Quantitative representation of microbial and human protein categories derived from proteomic analyses of salt-encrusted CBs and associated UP samples. The bars are ordered from the first to the last collection time point (left to right). A given time point has a specific number, either one or both sample types (CB, UP) may have been collected and profiled. Number gaps do not indicate missed samples. The colored segments in bars above the x-axis display abundances of microbial species relative to the entire profiled proteome per sample. Colored segments in bars below the x-axis display the abundances of distinct sets of human proteins known to be enriched in specific cells, such as neutrophils and eosinophils (NEI) and erythrocytes (ERY), are part of the complement system and acute phase response (CAP) or belong to the keratin group (KRT) as indicators of epithelial cell shedding into the urinary tract. Data on human protein category quantities are relative to total human protein content per sample.

### While proteomic data reveal higher temporal variability in microbial species profiles of mucoid biofilms compared to encrusted CBs, retention of urothelial reservoirs during catheter exchange is also evident

The UP and CB samples from P1, P2, P8 and P9 were collected over 2, 6, 5 and 6 months, respectively. In polymicrobial biofilms without salt encrustation, Enterobacteriaceae family members, *P. aeruginosa* and *E. faecalis* were most often identified. There were instances of dominance by a single species and cases where two or more species were similarly abundant. In P1 (Fig. 3) and P8 (Fig. S1, Suppl. Data), most bacterial organisms varied strongly in abundance from one timepoint to another, but dominant microbes were retained or recurred over the entire sampling time. An exception was the sample 71CB that was colonized by a *Stenotrophomonas maltophilia* strain. In P2 and P9 (Fig. 3), polymicrobial profiles revealed community shifts in CBs over time. On the other hand, the P1 and P9 profiles showed a pattern of microbial cohabitation reminiscent of encrusted biofilms: the persistent presence of *E. faecalis* in CBs shared by one or more Gram-negative uropathogens over a series of catheters. Sudden changes in biofilm composition shown for P1 and P9 are effects of antibiotic treatments and described later. Rare pathogens in CBs included *Bifidobacterium scardovii* (P2 and P8), *Brevundimonas* (P2), *Bordetella hinzii* (P2) and *Campylobacter curvus* (P8). Normally commensal organisms included *Lactobacillus* (P2) and *Veillonella parvula* (P8). A fungal pathogen, *Candida albicans,* was identified as at low-abundances in several samples, but incomplete lysis may underestimate the true abundance of this fungus in CBs.

**Fig. 3.**
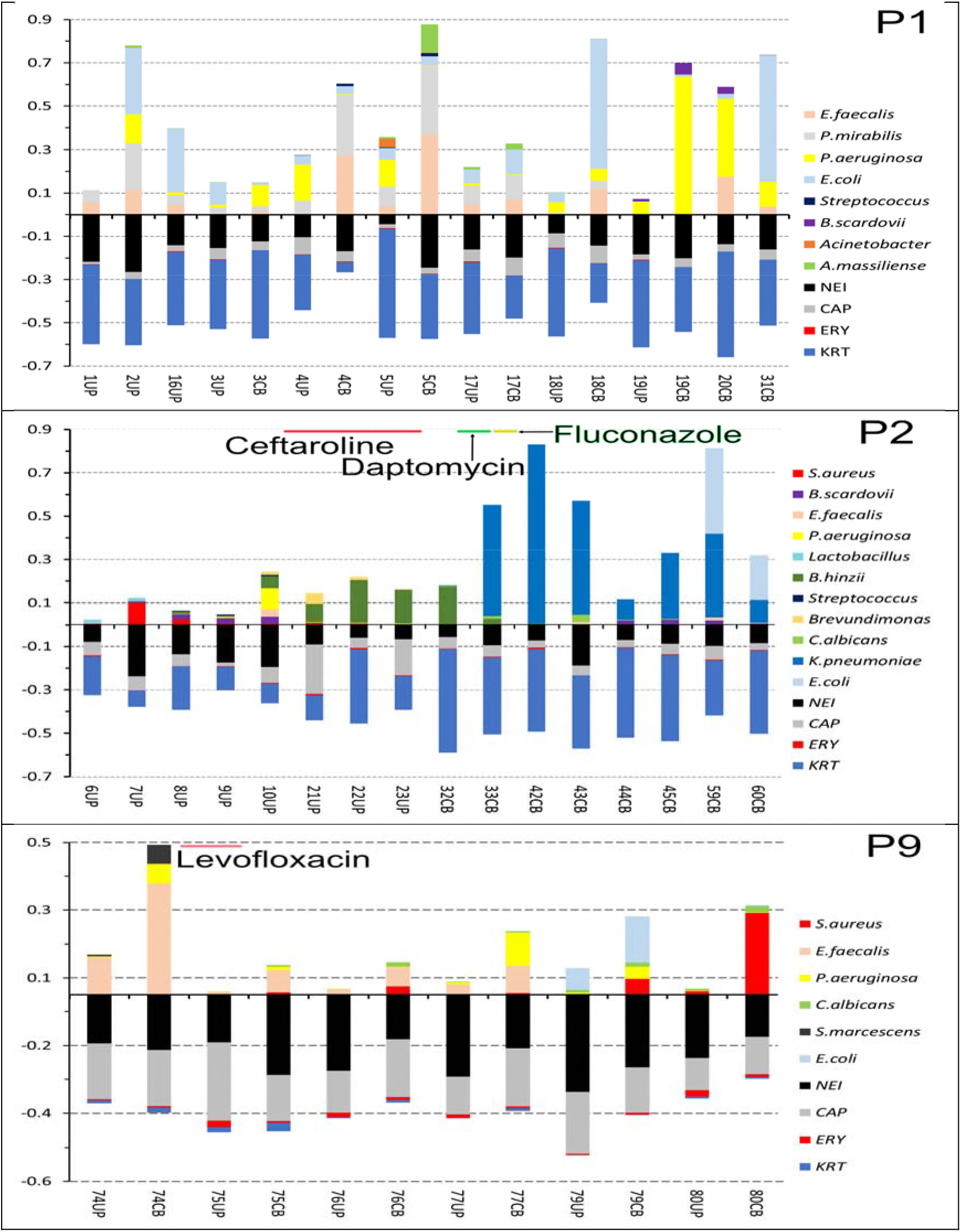
Quantitative representation of microbial and human protein categories derived from proteomic analyses of mucoid CBs and associated UP samples. The bars are ordered from the first to the last collection time point (left to right). Other information explaining the graphics is equivalent to that provided in the legend of Fig. 2. The time points and length of antibiotic treatment are indicated by lines above the name of the antibiotic in the context of the diagrams for P2 and P9.

The overall most prevalent pathogen in CBs was *E. faecalis.* Its contribution to the total microbial biomass varied considerably (Figs. 2 and 3). The observation suggests that *E. faecalis* colonizes quickly and fills a nutritional niche in polymicrobial biofilms. The species may have metabolic behaviors with mutual benefits for this and other species adapted to colonizing the catheterized human urinary tract. An *in vitro* study on enterococcal metabolites supporting *E. coli* growth was reported (26). However, *E. faecalis* can dominate as a colonizing microbe in long-term and short-term catheterization cases (P9, Fig. 3 and STC, Fig. S1, Suppl. Data, respectively). Bacteriuria diagnosed in short-term catheterized patients is typically caused by a single pathogen as reported (27) and re-displayed in Fig. S1, Suppl. Data. In conclusion, the long-term presence of catheters in the urinary tract facilitates the development of a polymicrobial biofilm with increased opportunities for less virulent and commensal bacteria to join a relatively stable community.

### Antibiotic drug treatment temporarily disrupts CBs with different long-term microbial colonization outcomes

Patient P7 was treated with oral sulfamethoxazol and trimethorprim drugs at timepoints 55UP/CB and 56UP (Fig. S1, Suppl. Data) for a wound infection. The antibiotic treatment cleared a *P. mirabilis* biofilm transiently. A bacterial reservoir recolonized upon completion of the treatment. Before and after treatment, the proteome of *P. mirabilis* revealed a high abundance of hemolysin (HpmA), a toxin not detected in *Proteus* strains that infected other patients. This supports the notion of strain-specific CAASB recurrence. *Haemophilus influenzae,* a minor component of the CBs also re-merged, but with a time gap of 3 months. A 10-day course of levofloxacin, taken by P9 to treat a wound infection, reduced the bacterial biomass from timepoint UP/CB74 to timepoint UP/CB75 (Fig. 3). While *Serratia marcescens* was eliminated, *E. faecalis* and *P. aeruginosa* persisted and a *C. albicans* strain newly colonized subsequently placed catheters. Levofloxacin does not have antifungal properties, and *E. faecalis* is known for high incidence of *parC* and *gyrA* mutations that confer fluoroquinolone resistance, as reported for UTI strains (28). Three antibiotic drug courses for a wound infection altered the structure of biofilms in patient P2. Intravenous cephalosporin treatments at timepoints corresponding to 10UP-23UP eliminated S. *aureus, E. faecalis, B. scardovii* and *P. aeruginosa* rapidly and *Brevundimonas* more slowly. The drug was ineffective in clearing *B. hinzii* (Fig. 3). When treatment was switched to daptomycin several days later, a lipopeptide antibiotic used to treat multidrug-resistant Gram-positive cocci, contributions of *B. hinzii* to the biofilm decreased (33CB), and a daptomycin-insensitive *K. pneumoniae* strain emerged and persisted in subsequent CBs. The antifungal drug fluconazole was administered to treat a fungal wound infection of P2 and did not have a lasting effect on the CB composition.

### Evidence of strong innate immune responses and embedding of human proteinaceous matter in catheter biofilm matrices

The absence of UTI symptoms in the context of bladder catheterization of patients is not indicative of subdued immune responses and lack of pyuria (7, 27). The data of our proteomic surveys in conjunction with catheter extract biomasses show that innate immune responses towards colonizing bacteria in CBs are chronic. Proteins enriched in activated neutrophils/eosinophils (NEI) and complement factors and other acute phase response proteins were abundant in many UP and CB samples. There were quantitative variations comparing specimens (Figs. 2 and 3), with a trend towards relative decreases of NEI quantities when KRT proteins were abundant (e.g., comparing P2 and P6 with P9). NEI quantities were low (< 15% of total human protein) when microbes were nearly absent in samples (e.g., 6UP, 47UP/CB and 55UP/CB; Figs. 2, 3 and S1). In previous work, we reported that cases of acute UTI and ASB associated with short-term catheterization have high NEI values, in contrast to absence of bacteriuria (12, 29). To gain confidence in the data assigning effector proteins to granulocytes rather than other immune cells, we determined average abundances of protein markers that are unique to immune cell types from all 121 datasets (Table 2). Among cell-specific surface markers, the granulocyte-specific CEACAM8 and the neutrophil-specific CD177 were far more abundant than CD3 (a T-cell marker), urothelial umbrella cell-specific uroplakins (e.g. uroplakin-2), CD19 (a B-cell and DC surface marker), integrin α-X and CD33 (both expressed on monocytes and macrophages). The quantification of red blood cell proteins (ERY) revealed microhematuria in many patients, while significant hematuria was rare (e.g., 50UP, P5, Fig. 2). In four patients (P1, P2, P7 and P8), KRT levels were high; two of the patients (P2, P7) were female. High KRT abundances in the absence of uroplakins support the notion that squamous epithelial cells exfoliate from the urethral mucosa in the infected urinary tract and are embedded in the extracellular CB matrix.

**Table 2.** Rank abundance of cell specific markers and neutrophil or eosinophil-enriched effectors

| UniProt No. | Rank No. <sup>1</sup> | Protein name | Cell association |
| --- | --- | --- | --- |
| P05164 | 5 | Myeloperoxidase | highly abundant in activated neutrophils |
| P07911 | 18 | Uromodulin | secreted from kidney tubules into urinary tract |
| P11678 | 68 | Eosinophil peroxidase | highly abundant in activated eosinophils |
| P69905 | 72 | Hemoglobin subunit alpha | highly abundant in erythrocytes |
| P31997 | 269 | Carcinoembryonic antigen-related cell adhesion molecule 8 | cell specific marker for granulocytes (CEACAM8) |
| Q8N6Q3 | 668 | CD177 antigen | cell specific marker for neutrophils |
| P20702 | 1205 | Integrin alpha-X | T-cell, B-cell, NK-cell, DC, MAC, monocyte and granulocyte cell specific marker <sup>2</sup> |
| P16422 | 1474 | Epithelial cell adhesion molecule | epithelial cell specific marker |
| P09693 | 3268 | T-cell surface glycoprotein CD3 gamma chain | T-cell specific marker |
| O00526 | 6276 | Uroplakin-2 | specific marker for urothelial umbrella cells |
| P20138 | 8141 | Myeloid cell surface antigen CD33 | DC, MAC, granulocyte, monocyte and stem cell marker <sup>2</sup> |
| P15391 | 8188 | B-lymphocyte antigen CD19 | B cell, DC and stem cell marker |

<sup>1</sup> used to estimate the relative spectral count-based abundances based on the rank number of proteins averaged from all datasets; <sup>2</sup> DC: dendritic cells; MAC, macrophages; NK natural killer cells

### Cooperativity and competition among bacteria cohabitating the CB niche viewed from a proteomic perspective

Bivalent metal ions, especially iron, and heme are sequestered to restrict the growth of microbial pathogens invading the human body. Transferrin (TRF) binds iron in blood plasma. LTF and calprotectin sequester iron and zinc, respectively, upon their release from mucosal cells and leukocytes. Lipocalin-2 (LCN2) binds enterobactin, a siderophore produced by Gram-negative bacterial pathogens, especially Enterobacteriaceae. To overcome the starvation of iron and other bivalent metal ions, the bacteria produce iron/siderophore receptors, metal ion inner membrane transporters and siderophore biosynthesis systems. We detected TRF, LTF and LCN2 at high abundances in many UP and CB proteomes. It was of interest to identify potential cooperative and competitive behaviors among two or more bacterial species present in a biofilm from a series of timepoints. By focusing on *E. coli* and *P. mirabilis* we observed that there was an ample supply of putative and known bivalent metal ion transporters, receptors and iron storage proteins (Dataset S4, Suppl. Data). Most iron acquisition and siderophore synthesis systems varied in their expression levels, for the colonizing strains, from patient to patient. Even functionally ill-characterized proteins, e.g. the TonB-dependent iron receptor PMI0842 *(P. mirabilis),* were expressed at very high levels. This data supports the notion that a distinct strain of a bacterial species typically dominates and persist in a series of CBs and favors recurrence of a strain rather than the patient’s re-infection with a different strain. We noticed frequent *E. faecalis* and *P. mirabilis* cohabitation in samples from P4 and, adding *E. coli,* samples from P1. The summed abundance of two putative *E. faecalis* metal uptake systems, one with a ZnuA-like and the other with a Fe/B12-type domain in their lipoprotein subunits, correlated well with the summed abundance of subunits of a *P. mirabilis* yersiniabactin-type siderophore synthesis and uptake system over the entire series of UP/CB samples for P1 (Fig. 4). A yersiniabactin system expressed by *E. coli* did not show an equivalent abundance correlation, but the data were more difficult to interpret because overall proteome abundances of *E. coli* and *E. faecalis* did not align well (Fig. 4). A weaker abundance correlation between the *E. faecalis* metal ion transporters and the *P. mirabilis* yersiniabactin system was determined for the data series of P4 (R = 0.49). To summarize, we hypothesize that *E. faecalis* uses two transporters, EfaABC and EF3082-3085, to acquire iron via use of a yersiniabactin-like xenosiderophore. A rare Gram-positive bacterial pathogen, *Globicatella,* also abundantly expressed a ZnuA-like ABC transporter predicted to import iron or bivalent metal ions while cohabitating a series of CBs with *P. mirabilis* (P6). However, the *P. mirabilis* strain dominant in the P6 samples did not express the yersiniabactin-type iron/siderophore synthesis and uptake system. The ZnuA-like lipoprotein of the transporter *(Globicatella* sp. HMSC072A10) is annotated as REF2811_06775.

**Fig. 4.**
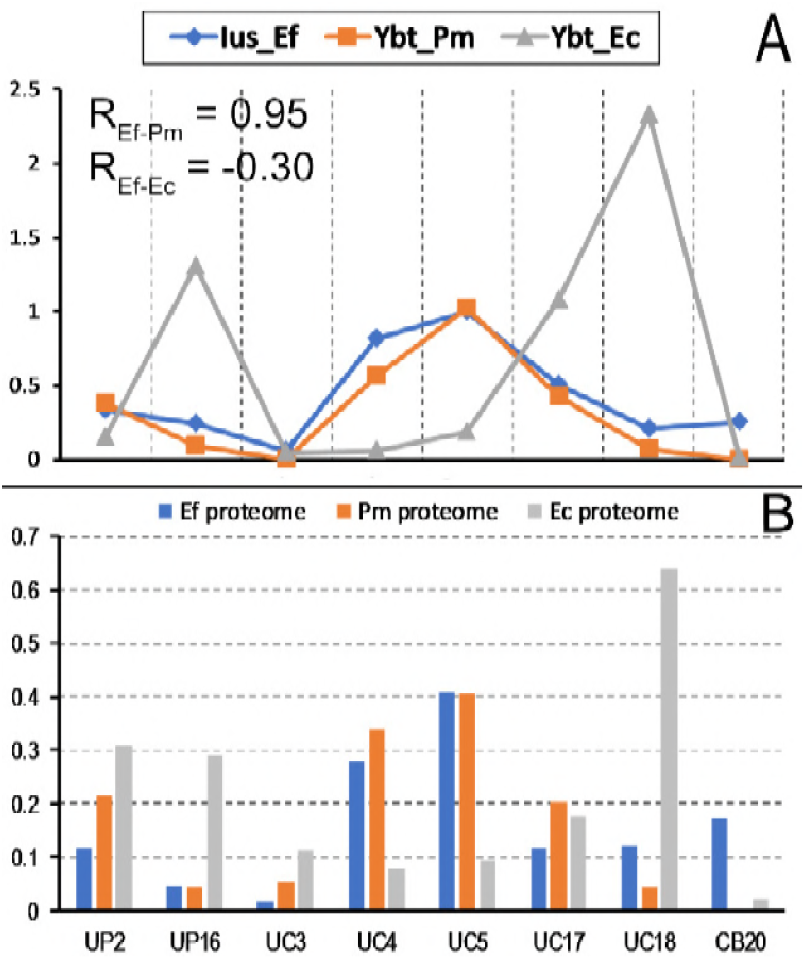
**(A)** Abundance correlations between *E. faecalis* metal ion uptake systems EfaABC and EF3082-3085 (denoted Ius_Ef) and *P. mirabilis/E. coli* yersiniabactin-type siderophore biosynthesis and uptake systems (denoted Ybt_Pm and Ybt_Ec, respectively, in the graphic) presented relative to total proteome at several sampling timepoint. The values for Ius_Ef are the summed abundances of proteins encoded by both transporters, EfaABC and EF3082-3085. On average, EfaABC was more than twice as abundant as EF3082-3085 in CB and UP samples. **(B)** For comparative purposes, the equivalent total bacterial species abundances based on proteomic data are provided for each timepoint. The term UC (in **B**) reflects instances where abundance values were combined from UP and CB data for a timepoint. The complete UP/CB sample series of patient P1 is presented.

Since bacteria compete with one another for nutrients, we assessed proteomic patterns indicative of competitive behaviors. Our focus was the expression of known and putative bacteriocins. The loss of contributions of *E. coli* and *P. mirabilis,* and to a lesser extent *E. faecalis,* to CBs of P1, previously dominated by such bacteria, occurred simultaneously with the surge of *P. aeruginosa* (Fig. 5). The *P. aeruginosa* strain expressed three factors tentatively linked to bacteriocin and cytotoxic functions: azurin (Azu), a type 6 secretion system (T6SS) and a pyoverdin biosynthesis (Pvd) system. Of these entities, only Azu was expressed in high abundance by a *P. aeruginosa* strain from another patient (72CB, P8). The abundance of Azu was highest at timepoint CB20 when *E. faecalis* had begun to recolonize the catheter surface, while the abundances of proteins associated with the T6SS and Pvd system were already in decline. Thus, our data favor a role of pyoverdin and/or the T6SS, with a yet undiscovered secreted cytotoxic substrate, in the ability of *P. aeruginosa* to out-compete most of the other catheter-colonizing bacteria. However, *B. scardovii* was unaffected by these conditions. As pointed out previously, a dominant *P. mirabilis* strain in UP and CB samples of P7 highly expressed a hemolysin (HmpA) which did not show any evidence of causing hemolysis in the urinary tract of this patient. HmpA was not detected in *P. mirabilis* proteomes profiled in the CBs of other patients. Only a single bacterial species, *H. influenzae,* seemed to be able to co-colonize the CBs of P7, with low or very low abundance for all nine timepoints. We hypothesize that HmpA, acting as a bacteriocin, restricts growth of other species in biofilms formed on the catheters from P7.

**Fig. 5.**
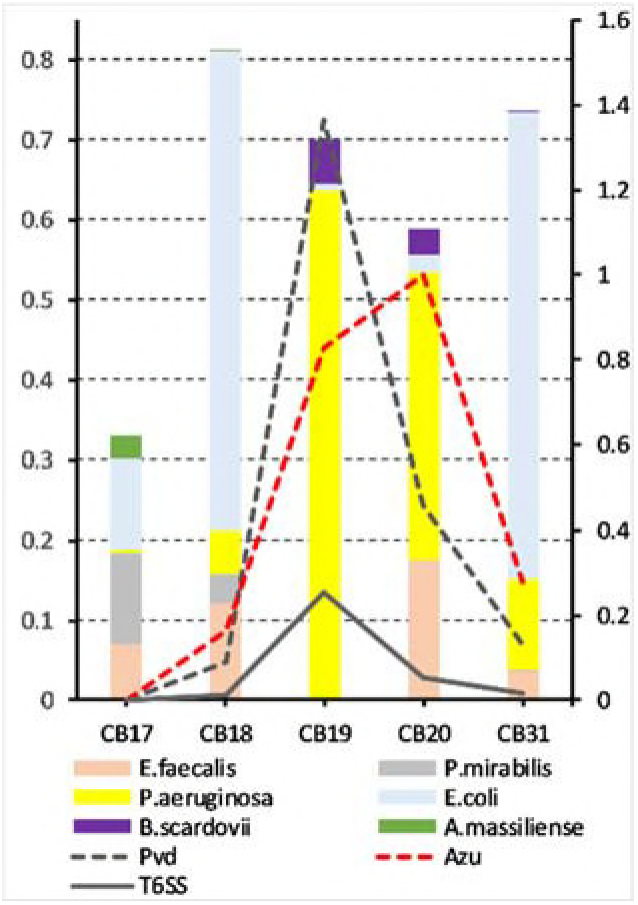
Bacterial colonization pattern for five consecutively collected catheters from patient P1 and abundance levels of *P. aeruginosa* systems potentially linked to bacteriocin functions and secretion. The bacterial abundance pattern, as described in the legend for Fig. 2 depicts representing relative contributions on the left y-axis (1 equals the entire proteome). The abundances of azurin (Azu) and the systems for type 6 secretion (T6SS) and pyoverdin synthesis and transport (Pvd) are represented in *%* of the total proteome on the right y-axis.

## Discussion

The objectives of our multi-year study on urethral catheter-associated biofilms were to gain insights into microbial population dynamics and host-pathogen interactions from the analysis of clinical specimens. Animal models do not mimic the polymicrobial nature of CBs and potential presence of fastidious bacteria that join such biofilms. Animal models do not reflect CB formation in serially replaced human catheters as the former rely on one-time catheter implantation. Low quantities of recoverable biological materials from murine studies are a limiting factor to study *in vivo* formed biofilms using less sensitive methods *(e.g.,* proteomics and metabolomics). Furthermore, differences in murine *vs.* human urothelial surface markers (16), in keratin expression patterns in urothelial and stratified squamous epithelial cells (30, 31) and in innate immune effectors such as DEFA1, a key antimicrobial peptide that is not produced by murine neutrophils, and TLRs (32) encourage the investigation of clinical samples with polymicrobial biofilms under challenge by the human immune system. Omics analyses from clinical samples may more accurately reflect host-pathogen interactions, characterize the extracellular biofilm matrix and determine the prevalence and roles of fastidious bacteria in CBs. Such analyses will potentially identify novel virulence factors as well as principles of inter-microbial species competition and cooperation in CBs. The outcomes will include new hypotheses on how to prevent the formation and enable the disruption of persistent CBs in the urinary tract. Recently developed human urothelial organoids (33), adapted to incorporate catheter plastics for microbial adhesion and biofilm growth, are a promising model to test the various hypotheses.

### Microbial diversity and long-term resilience of pathogens in CBs

By profiling catheter series longitudinally from neurogenic bladder patients, we found that microbial CBs persisted in all nine patients for up to ten catheter replacements, occurring over a 4-to 6-month time frame. The recurrence of CAUTI was previously reported for salt-encrusted biofilms dominated by *P. mirabilis* (3, 34). Indeed, our study also identified *P. mirabilis* as the dominant bacterial species in salt-encrusted CBs from three patients. Proteomic data support the notion that a single strain dominates the microbial community in a series of CBs from a distinct patient, enabled by survival and competitive growth advantages of that strain in the urinary tract during catheter replacements. The strain rapidly recolonizes a newly inserted sterile catheter. For instance, proteomic profiles of CBs from P7 revealed invariably high quantities of a hemolysin (HpmA). This cytotoxin was absent in CBs from P1, P4, P5 and P6, all of which were colonized by *P. mirabilis* strains. In contrast, CB profiles from P1 and P4 displayed high expression levels of a Ybt-like siderophore system, located in the PMI2596-PMI2605 gene cluster of strain HI4320 where it was first characterized (35). We cannot rule out the possibility that less abundant – recurrent or newly colonizing – *P. mirabilis* strains also contributed to CB profiles for some patients. Our data support the concepts that mucoid CBs also constitute recurrent biofilms and that the identified species are represented by dominants strains over a series of replaced catheters in a patient. For instance, only P1 and P2 had *E. coli* proteomes featuring a highly expressed a Ybt siderophore system (36), located in the C2178-C2187 gene cluster of strain UTI89. In contrast, P5 and P8 profiles, which also had evidence of long-term *E. coli* colonization, did not express a Ybt system. To provide another example arguing in favor of single strain dominance for various species in a CB series, the *P. aeruginosa* proteomes from CB18 to CB31 (P1) revealed highly abundant type 4 fimbriae components. Together with flagellins, type 4 fimbriae are in control of adhesion and mobility behaviors on mucosal surfaces (37). They were not confidently identified in CBs from P2, P8, P9 and early P1 samples with evidence of *P. aeruginosa* colonization. Finally, plasmid-derived *E. faecalis* proteins involved in resistance to neutrophil killing (e.g., PrgA, PrgB and Sea1) (38) were identified only in P1 and P4 samples; *E. faecalis* strains were detected in CBs from all patients. In conclusion, dominant strains of major pathogens tend to be responsible for re-infection of catheters in patients over a series of replaced catheters. Rare pathogens (e.g., *B. hinzii, A. massiliense, A. urinae, S. marcescens* and *Stenotrophomonas)* and a few species largely considered to be mucosal commensals (e.g., *V. parvula, B. scardovii* and *Lactobacillus)* also colonized CBs under study. In P2 and P5, rare pathogens emerged as substantial contributors to the CB microbial biomass (Figs. 2 and 3). These species have rarely been linked to CAUTI and UTI, and were not mentioned in reviews for the infections (1, 3, 4). We conclude that polymicrobial urethral and/or bladder reservoirs, dispersed from the CB surfaces followed by adhesion to mucosal surfaces, recolonize newly placed catheters in the form of groups. Bladder wash steps are not effective in preventing the recolonization. We do not rule out intra-urothelial quiescent life stages, as reported for *E. coli* and *K. pneumoniae* in the context of recurrent UTI (39, 40), as sources of pathogen recurrence on CBs. Bacteria that are retained in viable extracellular or intracellular reservoirs likely have growth advantages over microbes newly ascending from the urethral meatus. Our surveys suggest that CBs are dominated by bacteria, with only occasional presence of uropathogenic fungi. These CBs are more diverse than clinical cases of UTI and short-term catheterization.

### Antibiotic treatments transiently remove but ultimately fail to clear CBs

Data for three patients show that systemic antibiotic treatment may or may not have long-lasting effects on CB composition. While the bacterial biomass in CBs was reduced during the time of antibiotic treatment in two cases followed by the resurgence of previously identified species (P7 and P9), a third case revealed the elimination of susceptible bacteria during antibiotic treatment and emergence of pathogens resistant to the selected antibiotics. Robust re-colonization of catheters upon treatment completion was observed (P2). Exposure to cephalosporin resulted in outgrowth of a *B. hinzii* strain insensitive to this antibiotic. A few weeks later, exposure to daptomycin restricted the growth of *B. hinzii,* but allowed a *K. pneumoniae* strain to persist in subsequent biofilms in P2 (Fig. 3). We did not analyze the genomes of the *B. hinzii* and *K. pneumoniae* strains to identify any antibiotic resistance genes. However, our data highlight the threats associated with intermittent antibiotic treatment in the context of MDR development in CBs of long-term catheterized human subjects. Indeed, it was bacteremia in a patient catheterized during hip replacement that led to the emergence of the first β-lactam and carbapenem-resistant *K. pneumoniae* strain (41). Rational antibiotic treatment decisions for UTI or other symptoms in patients who use urethral catheters are very difficult to make. CBs are often too diverse for broad spectrum antibiotics to be effective, and treatments foster the exchange of genetic materials harboring antibiotic resistance genes. Biofilms are also naturally tolerant to antibiotics by limiting their penetration into deeper cell layers and via formation of quiescent persister cells (42).

### Evidence of chronic innate immune responses elicited by microbial CBs

Longitudinal proteomic analyses of CBs from all patients revealed evidence of neutrophil and eosinophil infiltration given that many of their cell-specific effectors were abundant in both CB and associated UP samples. To our knowledge, this data is the first demonstrating that the inflammatory responses to microbes present in CBs do not diminish over time in patients. We have evidence that the inflammation does not primarily result from mechanical irritation of catheter insertion, as also elaborate on in a study using a CAUTI animal model (22). We observed that neutrophil and eosinophil effectors had markedly lower abundances in cases where pathogens were undetectable (6UP, 47UP, 55UP). Our data are consistent with published data (43) that the complement system is also activated in the course of the immune response to CBs and that fibrinogen, as an extracellular matrix component, facilitates the formation of biofilms by pathogens such as *E. faecalis* (44) at the catheter surface. To our knowledge, we report for the first time that keratins are the most abundant group of host proteins in the CB matrix although the release of KRT varied among patients; *e.g.,* CBs of P9 had very low KRT content. We hypothesize that catheterization itself and CBs contribute to an inflammatory milieu promoting keratinization and mucosal cell death, thus leading to desquamation in the urethra and urothelial cell exfoliation in the bladder. The cell debris, rich in KRTs, is deposited in constantly renewing CBs. Both the presence of neutrophils and high abundance of KRTs were reported for *C. albicans* biofilms studied in a murine model of respiratory epithelial infection (45). In the context of biofilm formation, short-term catheterization (12) and long-term catheterization (7, 43) have been reported to result in pyuria, independent of whether patients report UTI symptoms or not.

### Siderophores, social traits and signaling molecules that influence pathogen invasion

Siderophores are microbial products linked to social traits, such as inter-microbial cooperation, competition and parasitism. Their synthesis and uptake affects bacterial strain fitness in a polymicrobial community (46). Their primary function is to acquire iron and other bivalent metals. These metal ions are sequestered by the host in mucosal tissues and body fluids to prevent by microbial growth. Microbes synthesize siderophores to capture iron and bivalent metal ions to support diverse oxidoreductive processes in cells. Recently, their roles in eliciting granulocyte infiltration and host invasion were identified in the context of *E. coli* and *K. pneumoniae* infections (47–49). Our methods to study CBs were suitable to examine siderophore function as a social trait and mediator of bacterial survival and urothelial invasion. We identified subunits of siderophore biosynthesis systems for nearly all Enterobacteriaceae members and *P. aeruginosa* in CBs. TonB-dependent outer membrane receptors for catecholate (enterobactin) and phenolate (yersiniabactin, pyoverdin, pyochelin) siderophores were co-expressed, supporting their roles in iron scavenging. Receptors for unknown substrates were also abundant, e.g. PMI0842 *(P. mirabilis)* in CBs from P7 and Smlt1426 *(Stenotrophomonas)* in 71CB. Siderophore biosynthesis gene clusters are not expressed at significant levels in iron-supplemented *in vitro* bacterial cultures, suggesting widespread iron and other bivalent metal starvation in the CB milieu. Based on their abundances in proteomic datasets, we inferred especially high levels of siderophore production in distinct CBs (derived from *E. coli, P. aeruginosa, P. mirabilis, K. pneumoniae and E. aerogenes* strains; *e.g.,* in P1 and P8). Based on reports on siderophores as promoters of inflammation and tissue invasion, we assessed whether the inferred high siderophore production levels correlated with high NEI and ERY scores in the UP and CB datasets. There was no evidence of marked increases in neutrophil and eosinophil effector abundances or of more extensive hematuria in these cases. However, the studies that linked siderophores to host invasion were acute infections (47–49), while CAUTI with serial catheterizations of patients is a chronic condition. The lack of cases with evidence of extensive hematuria was unexpected. Hematuria is commonly observed in the context of acute UTI and ASB (27, 29). We hypothesize that long-term urinary tract catheterization can alter the mucosal surface and perhaps cause squamous epithelial metaplasia, a pathology associated with the chronically inflamed urinary tract (50). This process may explain the reduced incidence of hematuria in the cohort under study here.

### Microbial cooperation in CBs

We conducted correlation analyses to assess cooperativity among pathogens co-colonizing a catheter surface in relationship to siderophore utilization. Data from CBs of P4 and P6 revealed dominant colonization by *P. mirabilis,* but also persistent cohabitation by one or two other species that potentially fill a niche: *E. faecalis* in samples of P4 and *Globicatella* and *P. stuartii* in samples of P6. Both *E. faecalis* and *Globicatella* are small Gram-positive cocci that share many metabolic pathways and may adapt to the more alkaline milieu of encrusted catheter biofilms. Cohabitation of *P. stuartii* and *P. mirabilis* was previously brought into context with higher urease activity and increased pathogenic potential (51). Our data showed that urease was produced only by *P. mirabilis.* The abovementioned Gram-positive cocci highly expressed binding proteins for bivalent metal ion transporters with ZnuA-like domains (EfaA for *E. faecalis;* REF2811_06775 for *Globicatella)* in the respective CBs, suggesting roles in cation acquisition to support Fe^2+^, Zn^2+^ and Mn^2+^ dependent metabolic pathways. EfaA has been identified as an ABC transporter subunit, and EfaA as an endocarditis surface antigen with adhesive properties (52), regulated by Mn^2+^ levels (53). Interestingly, the *Streptococcus pyogenes* ortholog MtsABC was shown to be required for iron transport (54). In our study, the *P. mirabilis* strains of P4 and P6 expressed multiple systems for high affinity iron and heme uptake. This included the heme uptake system Hmu (PMI1424-1430) and TonB-dependent receptors for iron/siderophores *(e.g.,* PMI0842 and IreA). In P4 CBs, *P. mirabilis* expressed the Ybt synthesis and uptake system (PMI2596-2605). We found that the sum of expression levels of the EfaABC transporter and another predicted *E. faecalis* iron transporter (EF_3082-3085) strongly correlated with the abundance of Ybt system subunits in the data series for P4 (Fig. 4). We hypothesize that there is cooperativity between *P. mirabilis* and *E. faecalis;* the Ybt produced by *P. mirabilis* makes bivalent metal ion available to *E. faecalis* via use of a xenosiderophore and at least one of its transmembrane ABC transporters. In further support, the Ybt system and the *E. faecalis* ABC transporters were also expressed in several CBs from P1, but did not correlate well in abundance perhaps due to additional cohabitating microbes. We assume that xenosiderophore use, together with the ability of *E. faecalis* strains to bind to fibrinogen (44) and urethral mucosal surfaces via the adhesin EbpA (EF_1091-1093) and an energy metabolism adaptable to different oxygenation levels allow this opportunistic pathogen to be so prevalent in CBs associated with long-term catheterization. The Ybt system was not expressed by the *P. mirabilis* strain dominating the P6 biofilms. Thus, we do not have data in support of iron/xenosiderophore uptake via orthologous, abundant ABC transporter of *Globicatella.*

### Microbial competition in CBs

Microbes compete in their host environments for nutrient resources. The biofilm lifestyle, as compared to planktonic cells, is thought to aggravate nutrient starvation, including bivalent metal ions. Another important element of direct competition is the production of bacteriocins, with bacteriocin producer strains (species) out-competing bacteria sensitive to such bacteriocins (55). We observed strong compositional changes in CBs over a short time in P1 and P8 where *P. aeruginosa* and *Stenotrophomonas* strains, respectively, surged in abundance. We detected three *P. aeruginosa* systems associated with cytotoxic functions as abundant in the respective proteomes (Azu, Pvd and T6SS) of P1; *E. coli, P. mirabilis* and *E. faecalis* strains were displaced in CBs over two timepoints. In the CB series for P7, a *P. mirabilis* strain was highly dominant, and other species were unable to cohabitate the catheters. The P7 proteomes, unlike other CB series with significant *P. mirabilis* contents, produced the hemolysin HpmA, an ortholog of the better characterized cytotoxin ShlA in S. *marcescens* (56). While functional roles of Azu, Pvd, T6SS and HpmA in the displacement of other bacteria from CBs need to be investigated further, we hypothesize that one or more of them contribute to the ability of producer strains to out-compete species sensitive to these toxins.

## Methods

### Ethics Statement

The Southwest Regional Wound Care Center (SRWCC) in Lubbock, Texas, and the J. Craig Venter Institute (JCVI, Rockville, Maryland) generated a human subject protocol and a study consent form (#56-RW-022), which were approved by the Western Institutional Review Board (WIRB) in Olympia, Washington, and by JCVI’s IRB in 2013. All human subjects were adults and provided written consent. Specimens were collected firsthand for the purpose of this study. There was a medical need to replace indwelling Foley catheters in patients for bladder management. Scientists at the main research site, JCVI, did not have access to data allowing patient identification. Patient metadata were encrypted. Electronic and printed medical records at the clinical site were retained for four years to facilitate integration of medical and molecular research data.

### Human subjects and study design

Nine human subjects who used indwelling Foley catheters were enrolled in a prospective study. They had irreversible spinal cord injuries (SCIs) and were diagnosed with neurogenic bladder syndrome. Catheter replacement was a medical need and part of routine patient care at SRWCC. Medical data included gender, ethnicity and the diagnosis of chronic wound infections and diabetes. The enrolled individuals provided up to 15 specimens (urethral catheters and/or urine) collected longitudinally in 1-to 4-week intervals depending on patients’ needs for medical care. Catheter specimens were cut into 1-inch pieces, placed in polypropylene tubes and stored at −20°C, minimizing external contamination by use of gloves and sterile razor blades. Urine samples were obtained from catheter bag ports swabbed clean with alcohol prior to collection. Urine aliquots of 20 to 50 ml were stored at −20°C. We do not exclude the possibility that infrequent draining of a catheter bag allowed some *ex vivo* microbial growth in the respective urine specimen. Thus, the quantitative ratios of microbes in urine sediments may not precisely reflect those from the originally voided samples. Tubes containing the specimens were kept frozen during transport and stored at −80°C until further use at the JCVI.

### Urine specimen extractions

The catheter materials were latex (from 8 patients) and silicone (from patient P7). The pH, color and turbidity of urine specimens and the crystallization of salts on the inside and outside of catheter tubes were noted. To obtain a urine pellet (UP) from a sample, an aliquot was thawed, adjusted to 20°C and, if acidic, neutralized with 1 M Tris-HCl (pH 8.1) to a pH range of 6.5 to 7.5 and centrifuged at 3,200 × g for 15 minutes. The urine supernatant (SU) was also collected. UP samples were washed with PBS and aliquoted for DNA, protein and metabolite extractions (10%, 45% and 45%, respectively). A urethral catheter piece was placed in a Falcon tube with 2 to 3 ml CHO buffer and equilibrated to 20°C. CHO buffer contained 100 mM sodium acetate (pH 5.5), 20 mM sodium meta-periodate and 300 mM NaCl. The suspension was vortexed for 1 min and incubated in a sonication water bath for 10 min. Biofilm materials detached via sonication and by scraping the plastic surface with a plastic spatula. Vortex, sonication and sample recovery steps were repeated once. The pH of the suspension was adjusted to 6.5 to 7.5 using 1 M Tris-HCl (pH 8.1) followed by centrifugation at 8,000 × g for 15 minutes to generate catheter biofilm supernatant (CB_sup_) and pellet (CB_pel_) fractions. The CB_sup_ fraction was concentrated to approximately 0.5 ml in an Ultrafree-4 filter unit (10 kDa MWCO) by centrifugation at 3,200 × g and exchanged into PBS. The CB_pel_ fraction was aliquoted as described for UP samples. Processed UP and CB samples were stored at −80°C until further use.

### Cell lysis and proteomic sample preparation

UP and CB_pel_ samples contained cellular and extracellular matter. In a 1:5 volume ratio, these samples were subjected to lysis in SED solution, consisting of 1% SDS (v/v), 5 mM EDTA and 50 mM DTT. Solubilized samples were sonicated in a Misonex 3000 sonication water bath (ten 30s on/off cycles at amplitude 6.5) followed by exposure to 90°C for 3 min and a 15 min-incubation step interrupted by several vortex steps at 20°C. These UP and CB_pel_ lysates were cleared by centrifugation at 13,100 × g for 10 minutes and analyzed by SDS-PAGE to visualize bands and estimate the total concentration of protein in lysates based on Coomassie Brilliant Blue-G250 staining intensities. CB_sup_ concentrates were also analyzed by SDS-PAGE. Aliquots of the extracts containing approximately 100 μg protein were processed using filter-aided sample preparation (FASP) in single-tube Vivacon filters with a 10 kDa MWCO (Sartorius AG, Germany). Sequencing-grade trypsin was used as the digestion enzyme according to methods reported previously (57). FASP-processed peptide mixtures were desalted using the Stage-Tip method (58), lyophilized and ready for LC-MS/MS proteomic analysis.

### Shotgun proteomics via LC-MS/MS

Desalted peptide mixtures derived from UP, CBsup and CB_pel_ samples were dissolved in 10 μl 0.1% formic acid (solvent A) and analyzed using one of two LC-MS/MS systems: (1) a high-resolution Q-Exactive mass spectrometer (MS) coupled to an Ultimate 3000-nano LC system; (2) a low-resolution LTQ-Velos Pro ion-trap mass spectrometer coupled to an Easy-nLC II system. Both systems (Thermo Scientific, San Jose, CA) were equipped with a FLEX nano-electrospray ion source at the LC-MS interface. Detailed LC-MS/MS analytic procedures were described for the Q-Exactive (29, 57) and the LTQ-Velos Pro (59) platforms previously. For Velos Pro analysis, peptides present in a sample were trapped on a C_18_ trap column (100 μm × 2 cm, 5 μm pore size, 120 Å) and separated on a PicoFrit C_18_ analytical column (75 μm × 15 cm, 3 μm pore size, 150 Å) at a flow rate of 200 nl/min. Starting with solvent A, a linear gradient from 10% to 30% solvent B (0.1% formic acid in acetonitrile) over 195 minutes was followed by a linear gradient from 30% to 80% solvent B over 20 min and re-equilibration with solvent A for 5 min. Following each sample, the columns were washed thrice using a 30-min solvent A to B linear gradient elution to avoid sample carry-over. For Q-Exactive analysis, LC was conducted as reported (29). Electrospray ionization was achieved by applying 2.0 kV distally via a liquid junction. Parallel to LC gradient elution, peptide ions were analyzed in a MS^1^ data-dependent mode to select ions for MS^2^ scans using the software application XCalibur *v2.2* (Thermo Scientific). The fragmentation mode was collision-activated dissociation with a normalized collision energy of 35% (LTQ-Velos Pro) and 27% (Q-Exactive). Dynamic exclusion was enabled. MS^2^ ion scans for the same MS^1^ *m/z* value were repeated once and then excluded from further analysis for 30s. Survey (MS^1^) scans ranged from a *m/z* range of 380 to 1,800 followed by MS^2^ scans for the selected precursor ions. The survey scans with the Q-Exactive were acquired at a resolution of 70,000 (m/Δm) with a *m/z* range from 250 to 1,800. Q-Exactive MS^2^ scans were performed at a resolution of 17,500. The ten most intense ions were fragmented in each cycle. Ions that were unassigned or had a charge of +1 were rejected from further analysis. Two or three technical LC-MS/MS replicates were run for all samples. Their raw MS files were combined for database searches.

### Computational proteomic data analysis

Raw LC-MS/MS files were analyzed using the Sequest HT algorithm integrated in the software tool Proteome Discoverer v1.4 (Thermo Scientific). Technical parameters and the database construction were described previously (29, 57). Only rank-1 peptides with a length of at least seven amino acids were considered. The FDR rates were estimated using the integrated Percolator tool with a (reverse sequence) decoy database. Protein hits identified with 1% FDR thresholds were accepted for data interpretation. The ‘protein grouping’ function was enabled to ensure that only one protein was reported when multiple proteins shared the same set of identified peptides. Initially, the database contents were reviewed protein sequence entries in the non-redundant Human UniProt database (release 2015-06; 20,195 sequences) and protein sequence entries for proteomes (in UniProt) pertaining to 23 bacterial species commonly detected in the UTI-infected human urinary tract and *Candida albicans.* Guided by genus assignments from the 16S rDNA operational taxonomic unit analysis, the proteomes for 28 bacterial species were added to evaluate their presence in the samples in a targeted manner. All species and genome/proteome identifiers are listed in Table S1 (Suppl. Data). In the context of quantitative proteomic analyses, databases were customized to represent those microbial species for which consensus results from initial LC-MS/MS searches and 16S rRNA data suggested detectable presence. Applied in a patient-specific manner, the approach allowed us to minimize incorrect peptide-spectral match (PSM) assignments of chemically indistinguishable tryptic peptides derived from protein orthologs of phylogenetically similar species. Inclusion of many proteins with high sequence identities *(i.e.,* the protein orthologs of bacterial species from the same genus and even genera from the same family) generates quantitative inaccuracies in shotgun proteomics experiments. We have empirically determined that the main problems with respect to urinary tract colonizing organisms are *Enterobacteriaceae.* Members of this family are prevalent and abundant in the infected urinary tract.

For quantitative assessments, the sum of PSMs, assigned to protein of origin, were counted in proportion to the total proteome for a dataset. Thus, normalized values for each identified protein were obtained. Using the Q-Exactive system, we had determined that quantitative analyses using PSM counts were in good agreement with a gold standard method for semi-quantitative proteomics, MS^1^ peak integration via the LFQ algorithm integrated in the MaxQuant software (13). The protein abundance values were logerithmized (log2) and then summed based on assignment to a microbial species or a human protein sub-group. Human protein grouping pertained to enriched presence of proteins in distinct cell types such as neutrophils and eosinophils (abbreviated NEI) and erythrocytes (abbreviated ERY) or to a distinct functional role in host defense such as the compliment system and acute phase response (abbreviated CAP) or the epithelial/epidermal barrier-forming cytokeratins (abbreviated KRT here). The Table S2 (Suppl. Data) lists the proteins assigned to these four categories.

### Metagenomic data analysis

The UP and CB_pel_ samples (5-25 μl volume) were thawed and resuspended in 300 μl TES buffer (20 mM Tris-HCl, pH 8.0, 2 mM EDTA, and 1.2% Triton X-100), We vortexed occasionally and incubated at 75°C for 10 min followed by cooling to 20°C. To this suspension, we added 60 μl chicken egg lysozyme (200 μg/ml), 5.5 μl mutanolysin (20 U/ml; Sigma Aldrich) and 5 μl linker RNase A, gently mixed and incubated for 60 min at 37°C. After addition of 100 μl 10% SDS and 42 μl proteinase K (20 mg/ml), bacteria were lysed overnight at 55°C. DNA was extracted by adding an equal volume of phenol: chloroform: isoamylalcohol (25:24:1; pH 6.6). The suspension was vortexed and centrifuged at 13,100 x g for 20 min. The aqueous phase was transferred to a clean sterile microcentrifuge tube. The residual sample was extracted again repeating the previous step using an equal volume of chloroform: isoamylalcohol (24:1) followed by centrifugation at 13,100 x g for 15 min. To the aqueous phase, we added 3 M sodium acetate (pH 5.2) at one tenth of the volume. DNA was precipitated by adding an equal volume of ice-cold isopropanol. Incubations at −80°C for 30 min or at −20°C overnight followed. Precipitated DNA was centrifuged at 13,000 x g for 10 min and washed with 80% ethanol. The centrifugation step repeated. After air drying, the DNA pellet was resuspended in TE buffer (20 mM Tris-HCl, pH 8.0; 1 mM Na-EDTA) and stored at −20°C. We described the preparation of the DNA library for the amplification of V1-V3 regions of 16S rDNA bacterial genes and the MiSeq (Illumina) sequencing method previously (60). We employed the UPARSE pipeline for the phylogenetic analysis (61). Operational taxonomic units (OTUs) were generated *de novo* from raw sequence reads using default parameters in UPARSE, the Wang classifier and bootstrapping using 100 iterations. Taxonomies were assigned to the OTUs with Mothur applying the SILVA 16S rRNA database version 123 as the reference database (62). Unbiased, metadata-independent filtering was used at each level of the taxonomy by eliminating samples with less than 2000 reads and OTUs present in less than ten samples. Filtered data were analyzed based on relative contributions of microbial genera in a distinct sample.

### Microbial cultures

Bacteria derived from CB_pel_ and UP samples stored at −80°C were cultured on blood and MacConkey agar plates to isolate *E. faecalis* and *P. mirabilis* strains, respectively. Single colonies were picked, Gram-stained and microscopically viewed. A culture stock derived from a single bacterial colony was generated and analyzed to confirm the identity of the suspected species. The culture stock was inoculated to generate a fresh pre-culture in 5 ml brain heart infusion (BHI) broth. A 50 μl inoculum was used to grow cells in a 20 to 30 ml BHI broth culture. Bacterial cell pellets were grown to stationary phase (12-15 hours) in a shaker at 880 rpm set at 37°C. The OD_600_ values reached for the *E. faecalis* and *P. mirabilis* aerobic suspension cultures were 0.4-0.6 and 0.9-1.2, respectively. Cell aggregates were observed for *E. faecalis* cultures. Bacterial cells were washed with a 20-fold volume of PBS and lysed in the SED solution as described for UP samples. The cycle of sonication, incubation at 20°C with occasional vortex steps and 3-min heat exposure at 95°C were applied twice to generate a lysate and recover its supernatant by high-speed centrifugation for 15 min.

## Acknowledgements

This work was supported by the National Institutes of Health grant R01GM103598 titled “Urethral catheter-associated polybacterial biofilm formation and dispersal”. The funder had no role in study design, data collection and interpretation, or decisions to submit the work for publication. We thank the Ruggles Family Foundation for the support in acquiring the Q-Exactive mass spectrometer.

